# ohun: an R package for diagnosing and optimizing automatic sound event detection

**DOI:** 10.1101/2022.12.13.520253

**Authors:** Marcelo Araya-Salas, Grace Smith-Vidaurre, Gloriana Chaverri, Juan C. Brenes, Fabiola Chirino, Jorge Elizondo-Calvo, Alejandro Rico-Guevara

## Abstract

Animal acoustic signals are widely used in diverse research areas due to the relative ease with which sounds can be registered across a wide range of taxonomic groups and research settings. However, bioacoustics research can quickly generate large data sets, which might prove challenging to analyze promptly. Although many tools are available for the automated detection of sounds, choosing the right approach can be difficult only a few tools provide a framework for evaluating detection performance. Here we present *ohun*, an R package intended to facilitate automated sound detection. *ohun* provides functions to diagnose and optimize detection routines and compare performance among different detection approaches. The package uses reference annotations containing the time position of target sounds in a training data set to evaluate detection routines performance using common signal detection theory indices. This can be done both with routine outputs imported from other software and detections run within the package. The package also provides functions to organize acoustic data sets in a format amenable to detection analyses. ohun also includes energy-based and template-based detection methods, two commonly used automatic approaches in bioacoustic research. We show how ohun automatically can be used to detect vocal signals with case studies of adult male zebra finch (*Taenopygia gutata*) songs and Spix’s disc-winged bat (*Thyroptera tricolor*) ultrasonic social calls. We also include examples of how to evaluate the detection performance of ohun and external software. Finally, we provide some general suggestions to improve detection performance.

## Introduction

Animal acoustic signals are widely used to address a variety of questions in highly diverse areas, ranging from neurobiology (Burgdorf *et al*. 2011; Schöneich 2020) to taxonomy (Köhler *et al*. 2017; Gwee *et al*. 2019), community ecology (Zsebők *et al*. 2021; Tiwari & Diwakar 2022), and evolutionary biology (Medina-García *et al*. 2015; Odom *et al*. 2021). The profuse usage of animal sounds in research relates to the fact that they can be easily collected using non-invasive methods. In addition, animal sounds can be obtained in various natural and unnatural settings, with equipment that has become increasingly inexpensive and broadly accessible (Blumstein *et al*. 2011; Sugai *et al*. 2019). Online repositories have also facilitated the study of these communication signals at larger taxonomic and geographic scales. However, adopting bioacoustic approaches may also imply large amounts of data (i.e., lots of recordings), which can be challenging to analyze manually (Gibb *et al*. 2019). As a result, a growing number of computational tools for automatically-detected animal sounds is increasingly available (reviewed by Stowell 2022), reflecting the need for better and more efficient automated approaches (Gibb *et al*. 2019).

Most available tools for the automatic detection of acoustic events are free software accessible to a wider range of users and scientific questions. However, this diversity of automated detection tools also posits a challenge, as it can be difficult to navigate (Stowell, 2022). In this regard, using standard approaches for evaluating the performance of automatic detection tools might prove helpful in informing researchers’ decisions about which method better fits a given question and study system (Knight *et al*. 2017). The performance of automated sound event detection routines has typically been evaluated using standard indices from signal detection theory (Knight *et al*. 2017; Balantic & Donovan 2020). In its basic form, performance is assessed by comparing the output of a detection routine against a ‘gold-standard’ reference in which all the target sounds have been annotated (hereafter called ‘reference annotation’). This comparison facilitates quantifying the number of sounds detected correctly (true positives), wrongly (false positives), and missed (false negatives), as well as additional metrics derived from these indices (e.g., recall, precision).

Nevertheless, the fact that sound events are not evaluated as discrete classification units but are instead embedded within a continuous string of sound demands additional information to fully diagnose the temporal precision of the detection performance. This is particularly relevant if identifying the precise time position of sounds is needed, which is often required when the main goal is measuring the acoustic structure of sounds (Araya-Salas & Smith-Vidaurre 2017). In this line, several detection problems can be encountered; for instance, the same signal can be detected as several separated sounds, the inferred time position can be offset from the target signal position, or several sounds can be detected as one single signal. Therefore, tools containing metrics that account for these additional performance dimensions are valuable for properly diagnosing automatic sound event detection.

Here we present the new R package *ohun*. This package is intended to facilitate the automatic detection of sound events, providing functions to diagnose particular aspects of acoustic detection routines to simplify their optimization. The package uses reference annotations containing the time position of target sounds that, along with the corresponding sound files, serve as a training data set to evaluate the performance of detection routines. This can be done with routine outputs imported from other software and detection routines run within the *ohun* package. The package also provides a set of functions to explore acoustic data sets and organize them in an amenable format for detection analyses. In addition, it offers implementations of two automatic detection methods commonly used in bioacoustic analysis: energy-based detection and template- based detection (Mellinger & Clark 2000; Charif *et al*. 2010; Aide *et al*. 2013; Bioacoustics 2014; Hafner & Katz 2015). Here, we explain how to explore and format acoustic data sets and how sound event detection routines can be evaluated.In addition, we showcase the package usage with study cases on male Zebra-finch songs (*Taenopygia gutata*) and Spix’s disc-winged bat calls (*Thyroptera tricolor*), which correspond to different recording settings (i.e., lab and flight cages) and signal types (i.e., sonic mating sounds and ultrasonic social calls).

### Formatting acoustic data sets

The format and size of the acoustic data to be analyzed needs to be standardize to avoid downstream errors and to inform expectations for computational time performance. Several functions in *ohun* can facilitate double-checking the format of acoustic data sets prior to automatic detection. The function feature_acoustic_data prints a summary of the duration, size, and format of all the recordings in a folder. Here we explore the acoustic data set of zebra finch’s songs (Supporting Information):

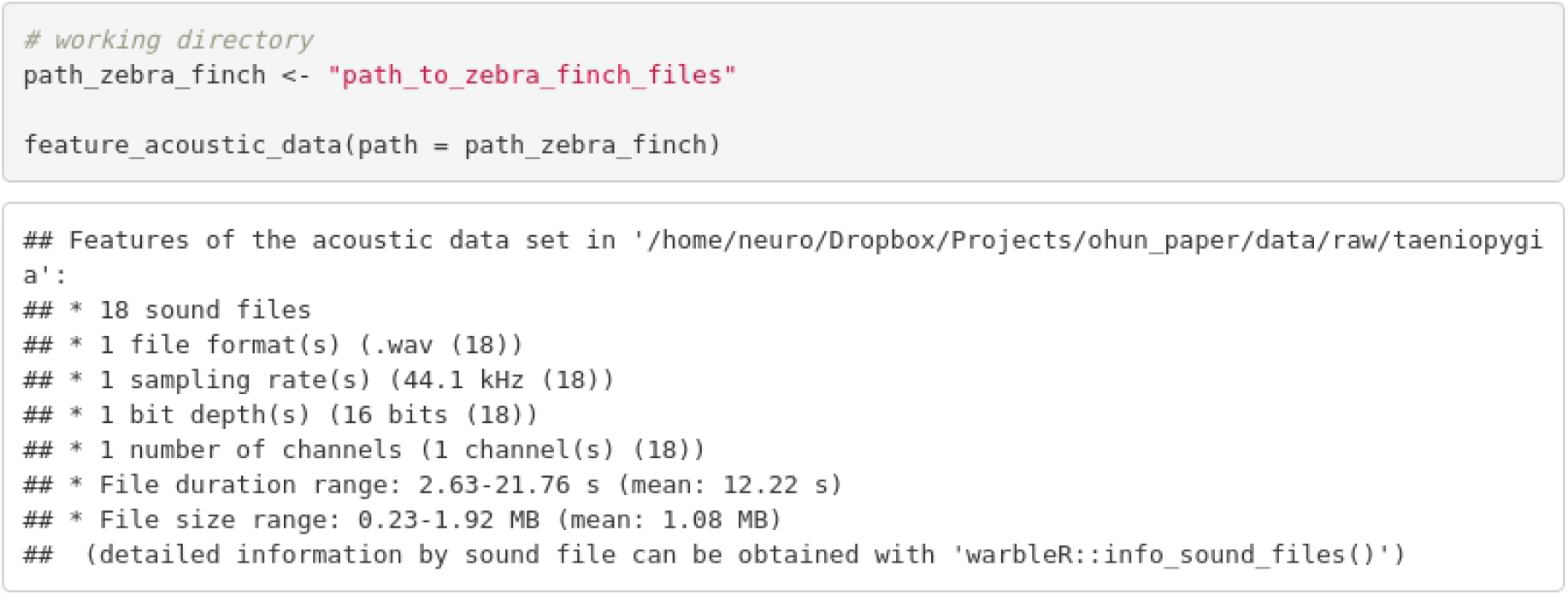

In this case, all recordings have the same format (.wav files, 44.1 kHz sampling rate, 16-bit resolution, and a single channel). We can also check the files’ duration and size. Format information is important as some tuning parameters of detection routines can behave differently depending on file format (*e.g*., time window size can be affected by sampling rate) or simply because some software might only work on specific sound file formats. In addition, long sound files could be difficult to analyze on some computers and might have to be split into shorter clips. In the latter case, the function split_acoustic_data can be used to produce those clips:

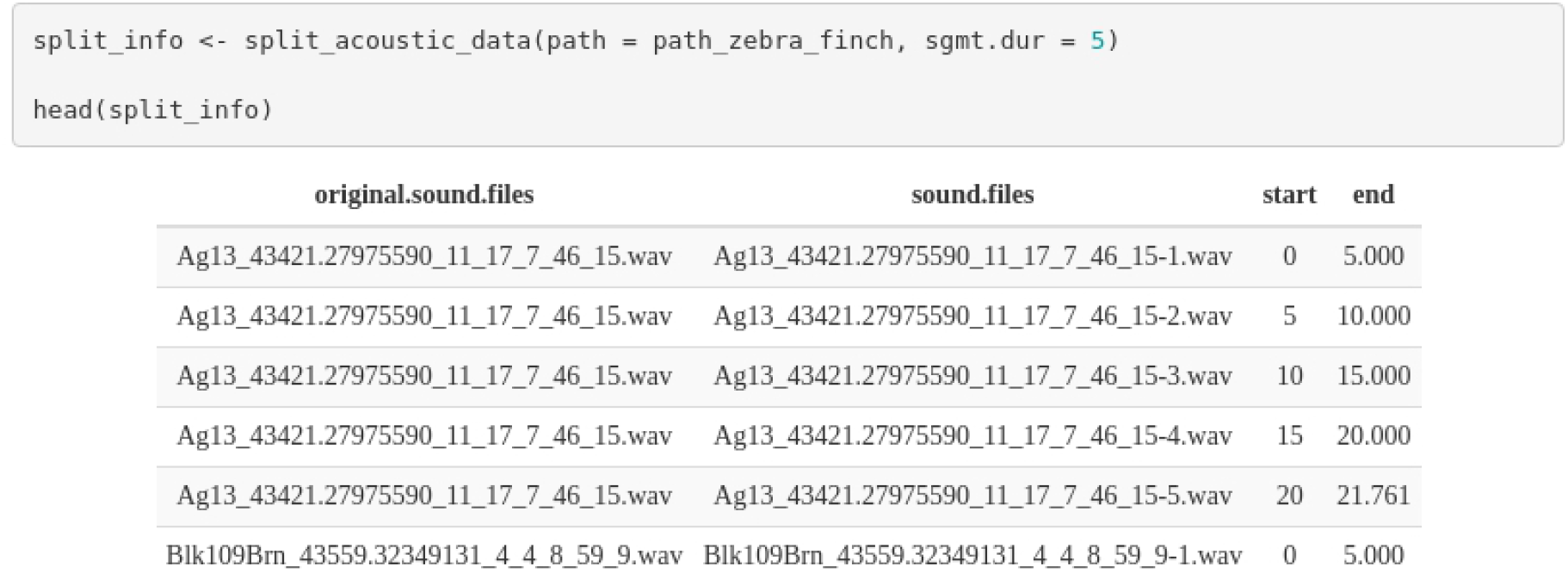

The output shows the time segments in the original sound files to which the clips belong. If an annotation table is supplied (argument ‘X’), the function will adjust the annotations, so they refer to the position of the sounds in the clips. This can be helpful when reference tables have been annotated on the original long sound files.

Annotations can also be explored using the function feature_reference, which returns the mean and range of signal duration and gap duration (e.g., time intervals between selections), bottom and top frequency, and the number of annotations by sound file. If the path to the sound files is supplied, then the duty cycle (i.e., the fraction of a sound file corresponding to target sounds) and peak amplitude (i.e., the highest amplitude in a detection) are also returned:

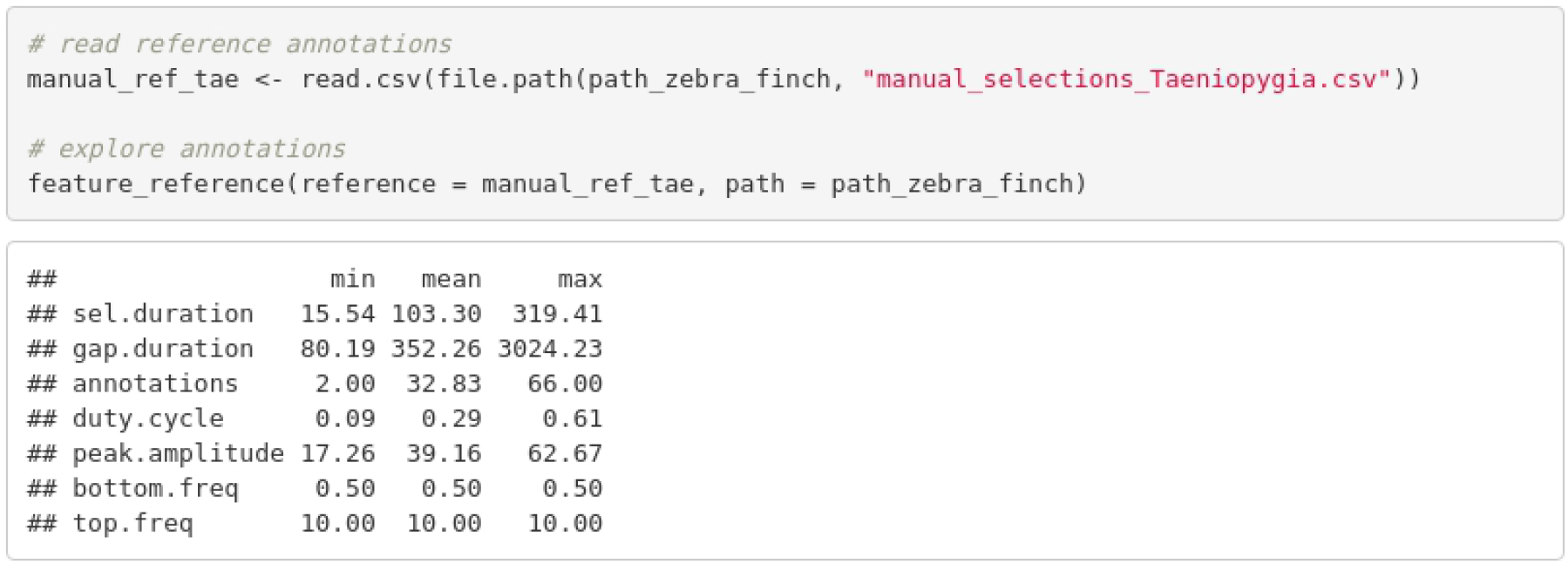

### Diagnosing detection performance

The *ohun* package uses signal detection theory indices to evaluate detection performance. Signal detection theory deals with the process of recovering sounds (*i.e*., target sounds) from background noise –not necessarily acoustic noise– and it is widely used for optimizing this decision-making process in the presence of uncertainty (Hossin & Sulaiman 2015). During a detection routine, the detected ‘items’ can be classified into four classes: true positives (TPs, target sounds correctly identified as signal), false positives (FPs, noise incorrectly identified as ‘signal’), false negatives (FNs, sounds incorrectly identified as noise), and true negatives (TNs, background noise correctly identified as noise). However, TNs cannot always be easily defined in the context of sound event detection, as noise cannot always be partitioned into discrete units. Hence, the package makes use of TPs, FPs, and FNs to calculate three additional indices that can further assist with evaluating the performance of a detection routine and are widely used in sound event detection (Knight *et al*. 2017): recall (i.e., correct detections relative to total detections), precision (i.e., the proportion of target sounds that were correctly detected) and F1 score (combined recall and precision as the harmonic mean of these two, which provides a single value for evaluating performance, a.k.a. F-score, F-measure or Dice similarity coefficient).

A perfect detection will have no false positives or negatives, resulting in both recall and precision equal to 1. However, perfect detection cannot always be achieved. Therefore, some compromise between detecting most target sounds plus some noise and excluding noise but missing target sounds might be warranted. These indices provide a useful framework for diagnosing and optimizing the performance of a detection routine. Researchers can identify an appropriate balance between these two extremes by the relative costs of missing sounds and mistaking noise for target sounds in the context of their specific study goals.

*ohun* offers tools to evaluate the performance of sound event detection methods based on the indices described above. To accomplish this, annotations derived from a detection routine are compared against a reference table containing the time position of all target sounds in the sound files. For instance, the following code evaluates a routine run in Raven Pro 1.6 (Charif *et al*. 2010) using the “band limited energy detector” option (minimum frequency: 0.8 kHz; maximum frequency: 22 kHz; minimum duration: 0.03968 s; maximum duration: 0.54989s; minimum separation: 0.02268 s) on a subset of the zebra finch recordings described below:

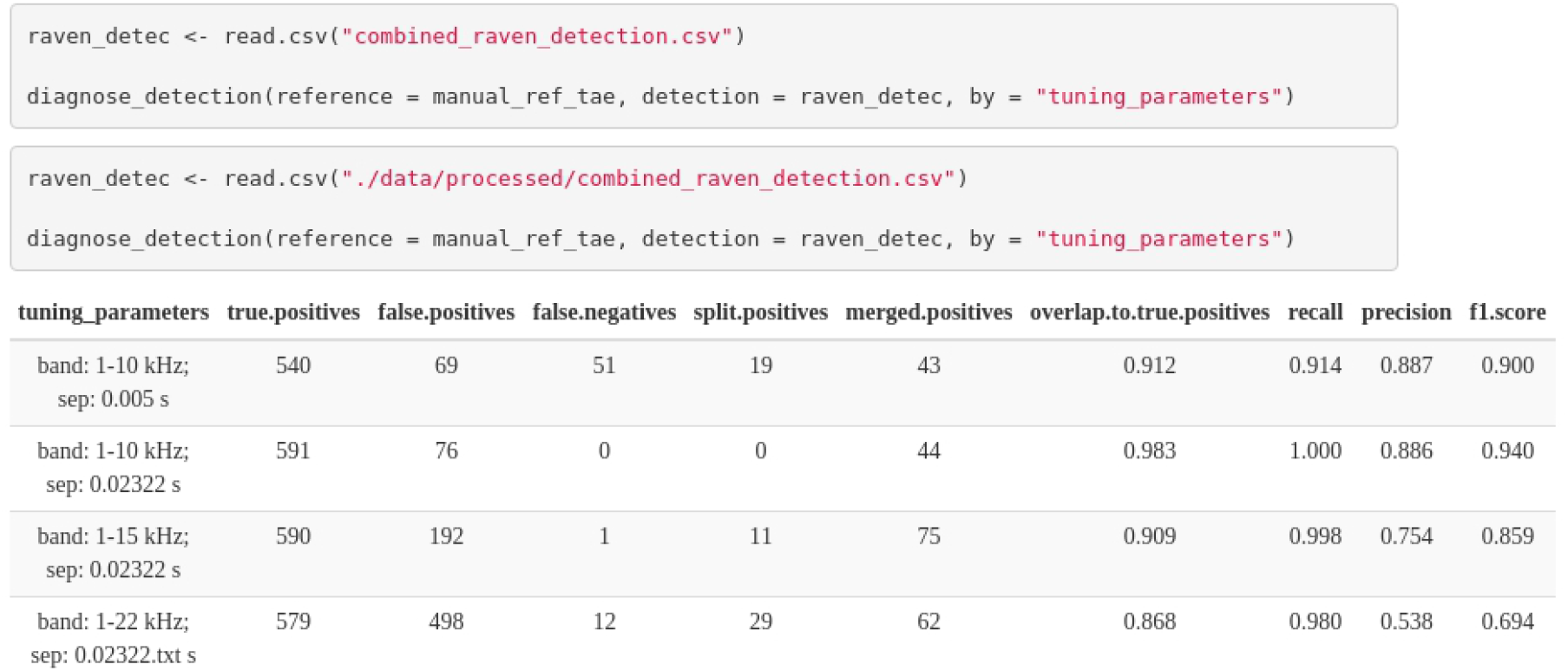

The function diagnose_detection uses bipartite matching graphs (Csardi & Nepusz 2006) to determine optimum matching of detected and reference events, ensuring that each reference sound is only be matched to a single detection (Lostanlen et al. 2019).

The output shows the indices described above, plus three additional metrics specific to sound event detection: ‘split positives’, ‘merge positives’, and ‘overlap to true positives’. ‘Split positives’ is the number of reference sounds overlapped by more than one detection, ‘merge positives’ is the number of detected sounds overlapping with another detection, and ‘overlap to true positives’ quantifies the mean overlap between detections and reference signal (1 means complete overlap). The function also allows detailing those indices by sound file as well as by additional categorical features. Here we show the first ten files detailed by the column ‘tunning_parameters’ which contains the combained detection parameter values used in Raven:

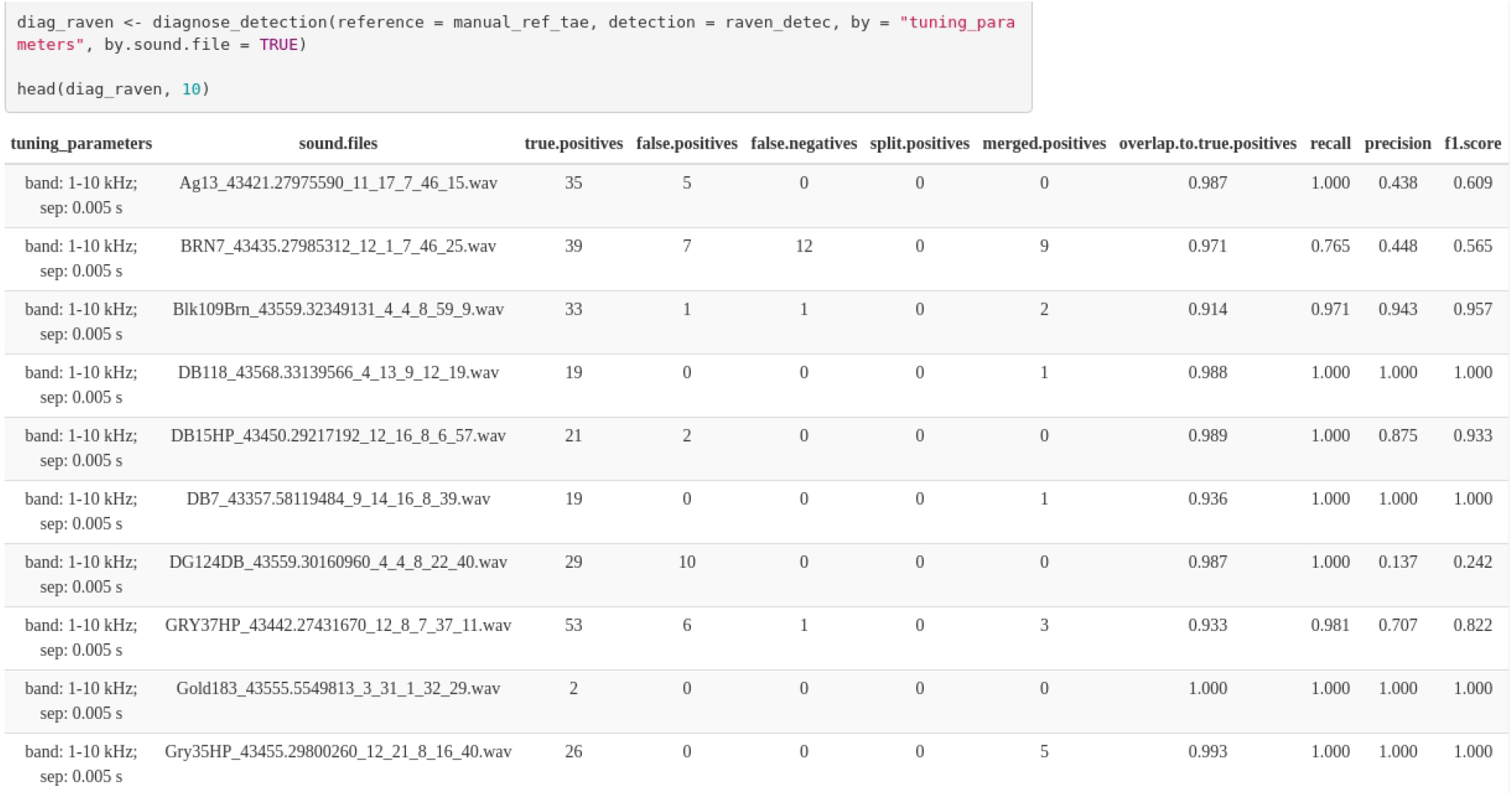

Diagnostics from routines utilizing different tuning parameters can serve to identify the parameter values that optimize detection. This process of evaluating different routines for detection optimization is incorporated into the two signal detection approaches provided natively by *ohun*, which we depict in the following section. Note that the detection with Raven Pro does not necessarily reflect the best performance of this software and has been included only as an example of evaluating detection from external sources rather than a direct comparison of performance between Raven Pro and *ohun*.

### Signal detection with *ohun*

The package offers two methods for automated signal detection: template-based and energy-based detection. These methods are better suited for stereotyped or good signal-to-noise ratio sounds, respectively. If the target sounds do not fit these requirements, more elaborate methods (i.e., machine/deep learning approaches) are warranted.

### Study cases

#### Template detection on ultrasonic social calls of Spix’s disc-winged bats

We recorded 30 individuals of Spix’s disc-winged bats (*Thyroptera tricolor*) at Baru Biological Station in southwestern Costa Rica in January 2020. Bats were captured at their roosting sites (furled leaves of Zingiberaceae plants). Each bat was released in a large flight cage (9 x 4 x 3 m) for 5 minutes, and their ultrasonic inquiry calls were recorded using a condenser microphone (CM16, Avisoft Bioacoustics, Glienike/Nordbahn, Germany) through an Avisoft UltraSoundGate 116Hm plugged into a laptop computer running Avisoft-Recorder software. Recordings were made at a sampling rate of 500 kHz and an amplitude resolution of 16 bits.

Recordings were manually annotated using Raven Pro 1.6 (Charif *et al*. 2010). Annotations were created by visual inspection of spectrograms, in which the start and end of sounds were determined by the location of the continuous traces of power spectral entropy of the target sounds. A total of 644 calls were annotated (~21 calls per recording). Annotations were made with a time window of 200 samples and 70% overlap and were then imported into R using the package Rraven (Araya-Salas 2020).

Inquiry calls of Spix’s disc-winged bats are structurally stereotyped (Chaverri *et al*. 2010). Most variation is found among individuals, although the basic form of a short, downward broadband frequency modulation is always shared (Fig. 1, Araya-Salas et al. 2021).

**Figure 1.**
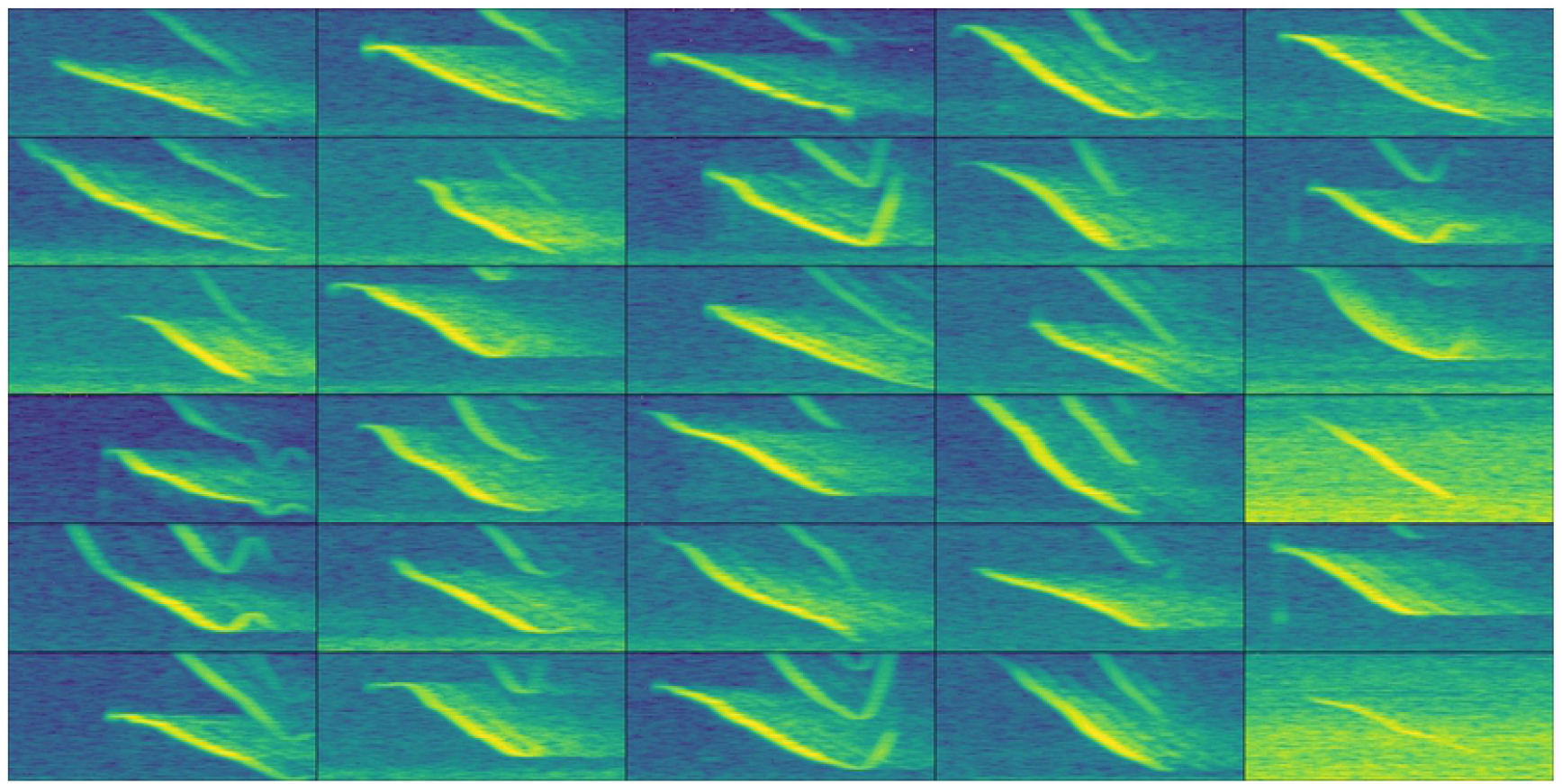
Example spectrograms of Spix’s disc-winged social calls for each of the 30 recordings used in the analysis. The highest signal-to-noise ratio call by sound file are shown. The time scale range is 71 ms and the frequency range 10-44 kHz.

A template-based detection is a useful approach when there are minimal structural differences in the target sounds (Knight *et al*. 2017; e.g., when signals are produced in a highly stereotyped manner, Balantic & Donovan 2020). It uses spectrographic crosscorrelation to find sounds resembling an example target sounds (i.e., template) across sound files, applying a correlation threshold to separate detections from background noise. We used this approach in *ohun* to detect inquiry calls. To do this, we tested the performance of three acoustic templates on a training subset of five sound files. First, we used the function get_templates to find several sounds representative of the variation in signal structure. This function measures several spectral features, which are summarized using Principal Component Analysis. The first two components are used to project the acoustic space. In this space, the function defines sub-spaces as equal-sized slices of a sphere centered at the centroid of the acoustic space. Templates are then selected as those closer to the centroid within sub-spaces, including the centroid for the entire acoustic space. The user needs to define the number of sub-spaces in which the acoustic space will be split.

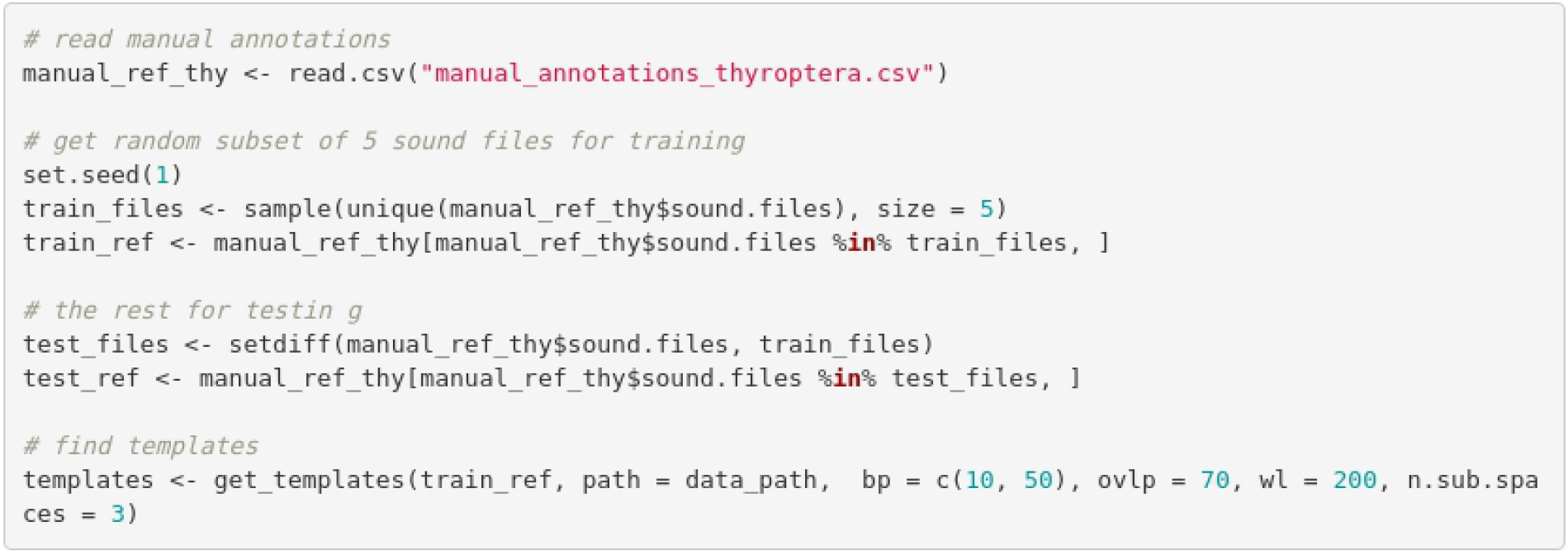

The output of the get_templates function includes an acoustic space plot (Fig. 2) in which the position of the sounds selected as templates is highlighted. Users can also provide their own acoustic space dimensions (argument ‘acoustic.space’). In the following code, we used the templates determined above for detecting bat social calls. The code iterates a template-based detection on the training data set across a range of correlation thresholds for each template, in order to find the combination of threshold and template with the best performance.

**Figure 2.**
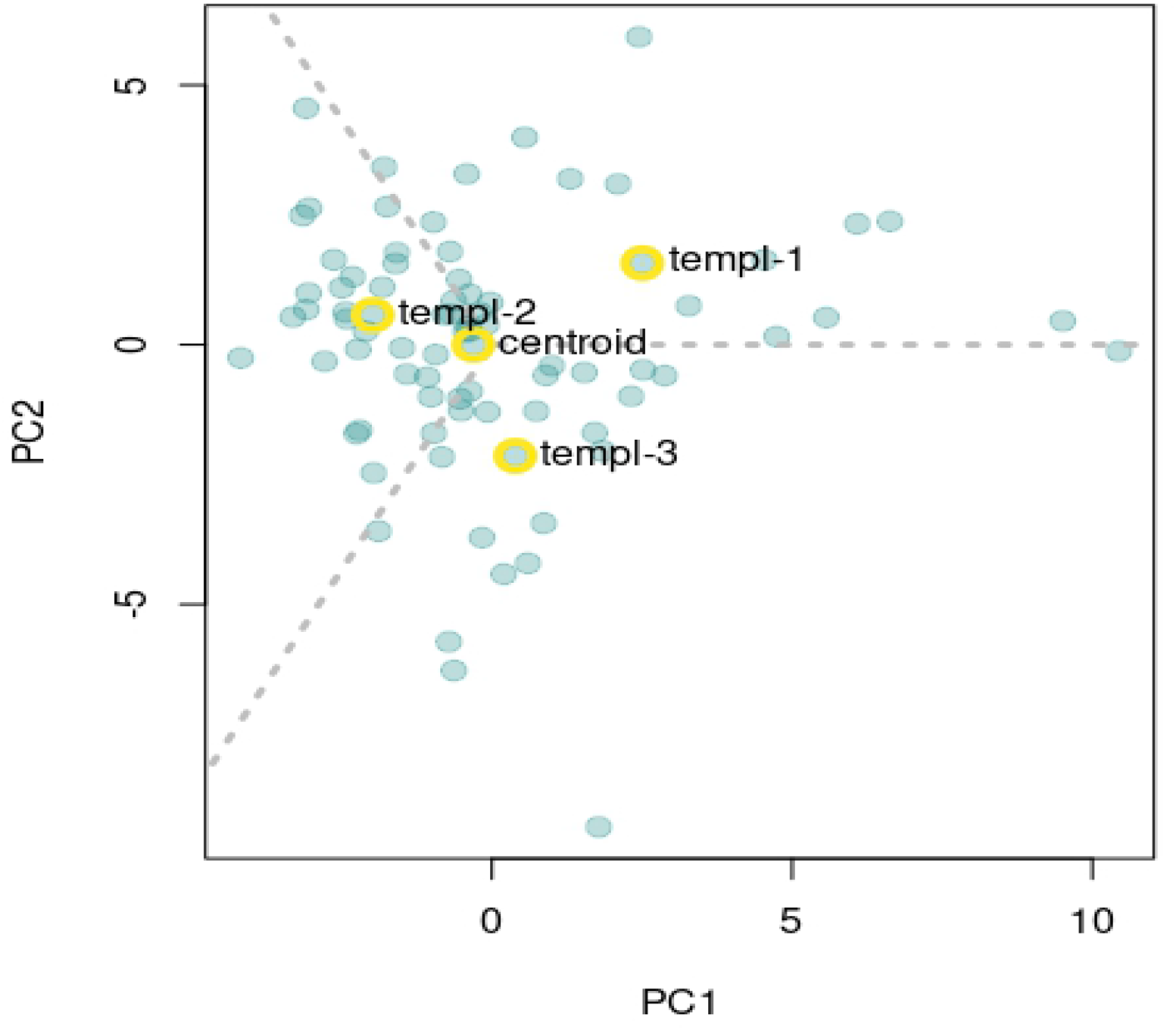
Acoustic space defined as the first two components of a Principal Component Analysis on spectrographic parameters. Templates are selected as those closer to the centroid within sub-spaces. Gray dashed lines delimit the region of sub-spaces. Yellow circles around points highlight the position of the signals selected as templates.

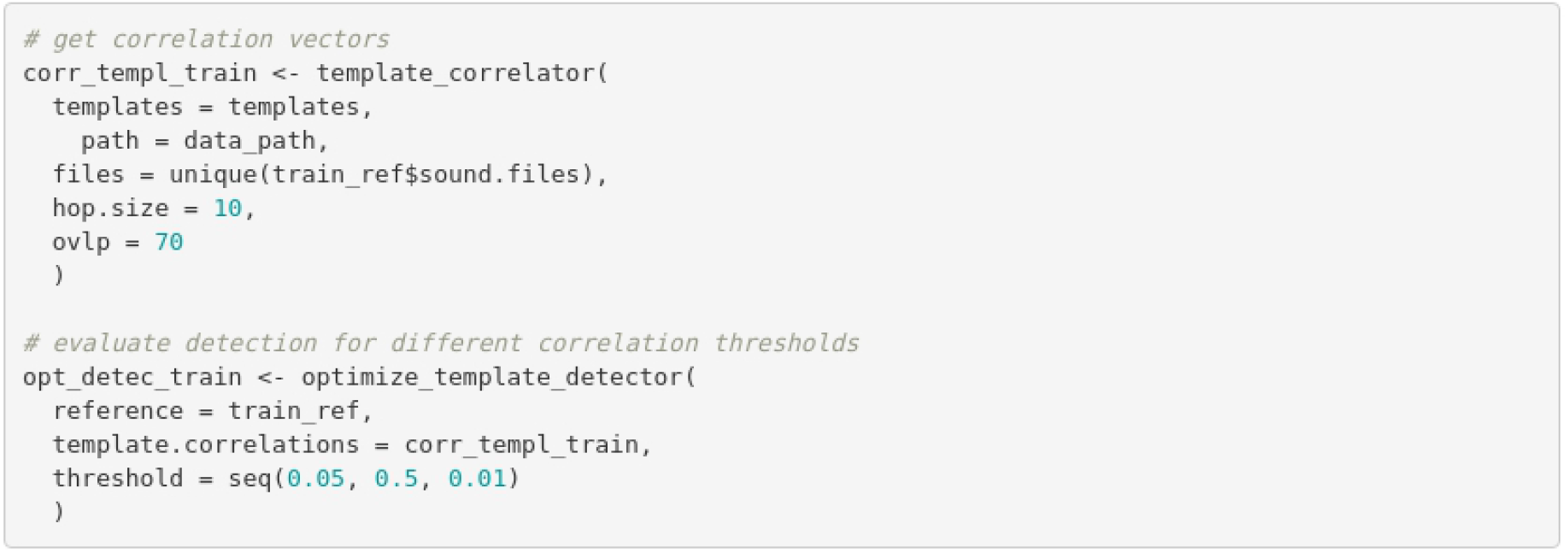

Note that the correlation vectors are estimated first (i.e., vectors of correlation values across sound files, template_correlator), and then the correlation thresholds are optimized on these vectors (optimize_template_detector). The output of optimize_template_detector contains the detection performance indices for each combination of templates and thresholds. Table 1 shows the two highest performance runs for each template.

**Table 1.**
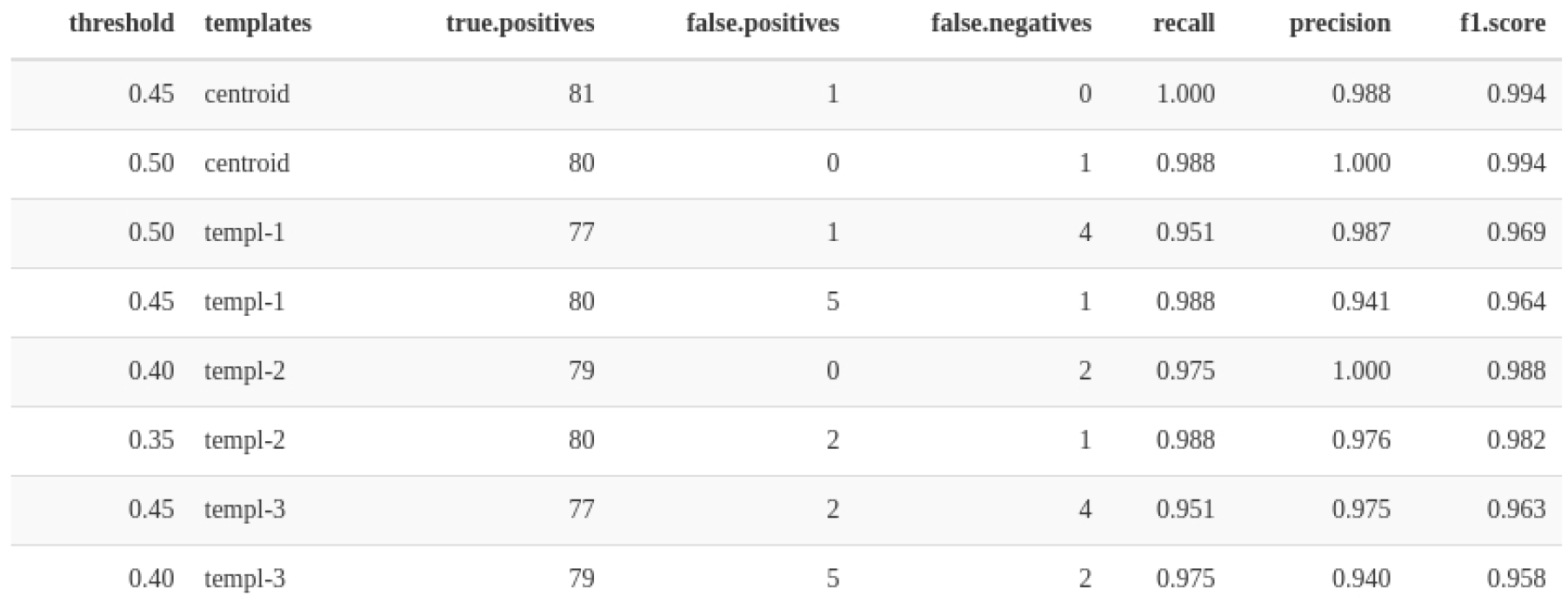
Performance diagnostic of template-based detections using four templates across several threshold values. Only the two highest performance iterations for each template are shown.

We can explore the performance of each template in more detail by looking at the change in F1 score across thresholds (Fig. 3).

**Figure 3.**
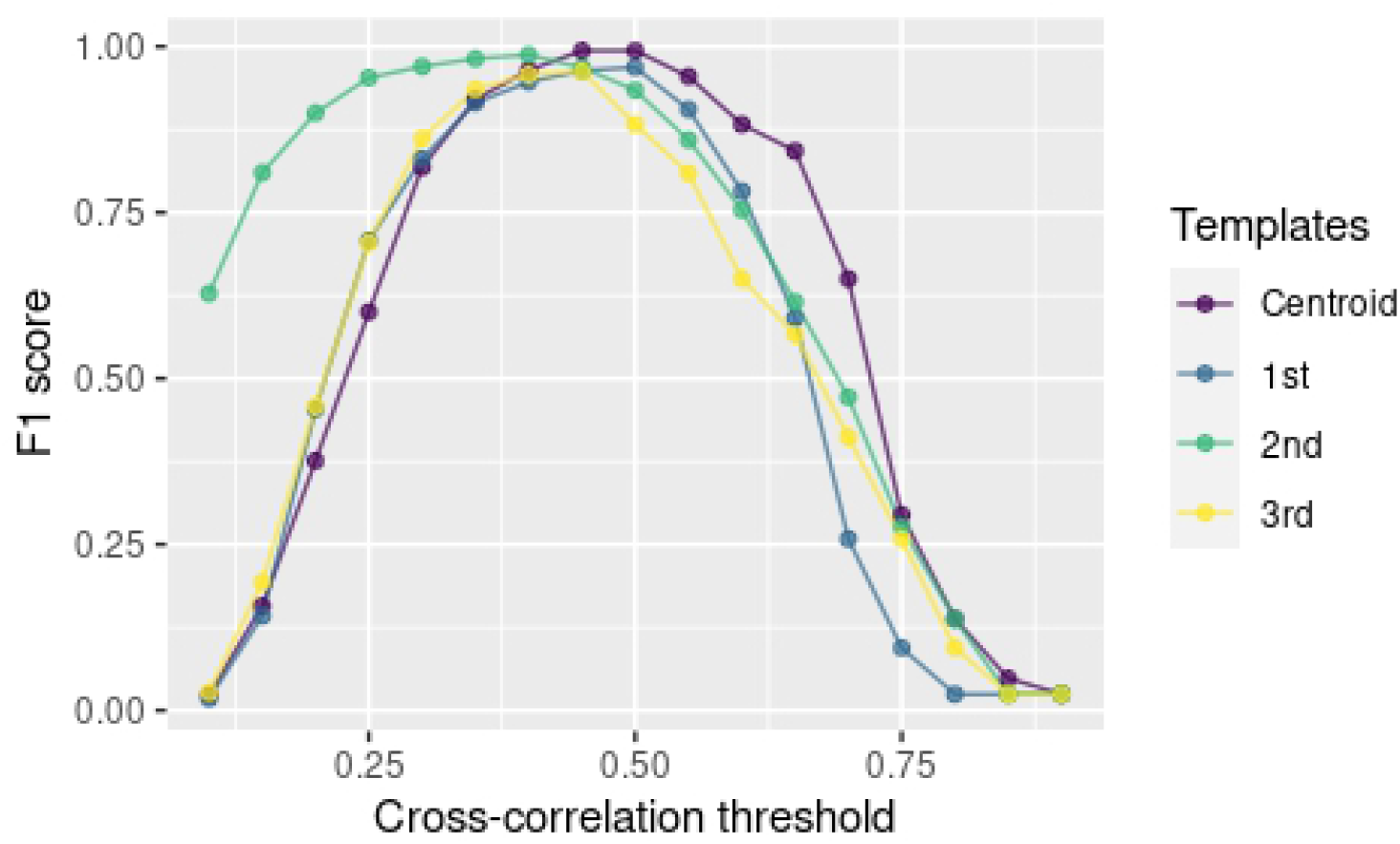
Changes in F1 score across the range of cross-correlation threshold values for four sound templates.

In this example, the “centroid” template produced the best performance (although not drastically different from other templates; Table 1; Fig. 3). Hence, we will use this template for detecting calls on the rest of the data. The following code extracts this template from the reference annotation table and uses it to find inquiry calls on the testing data set:

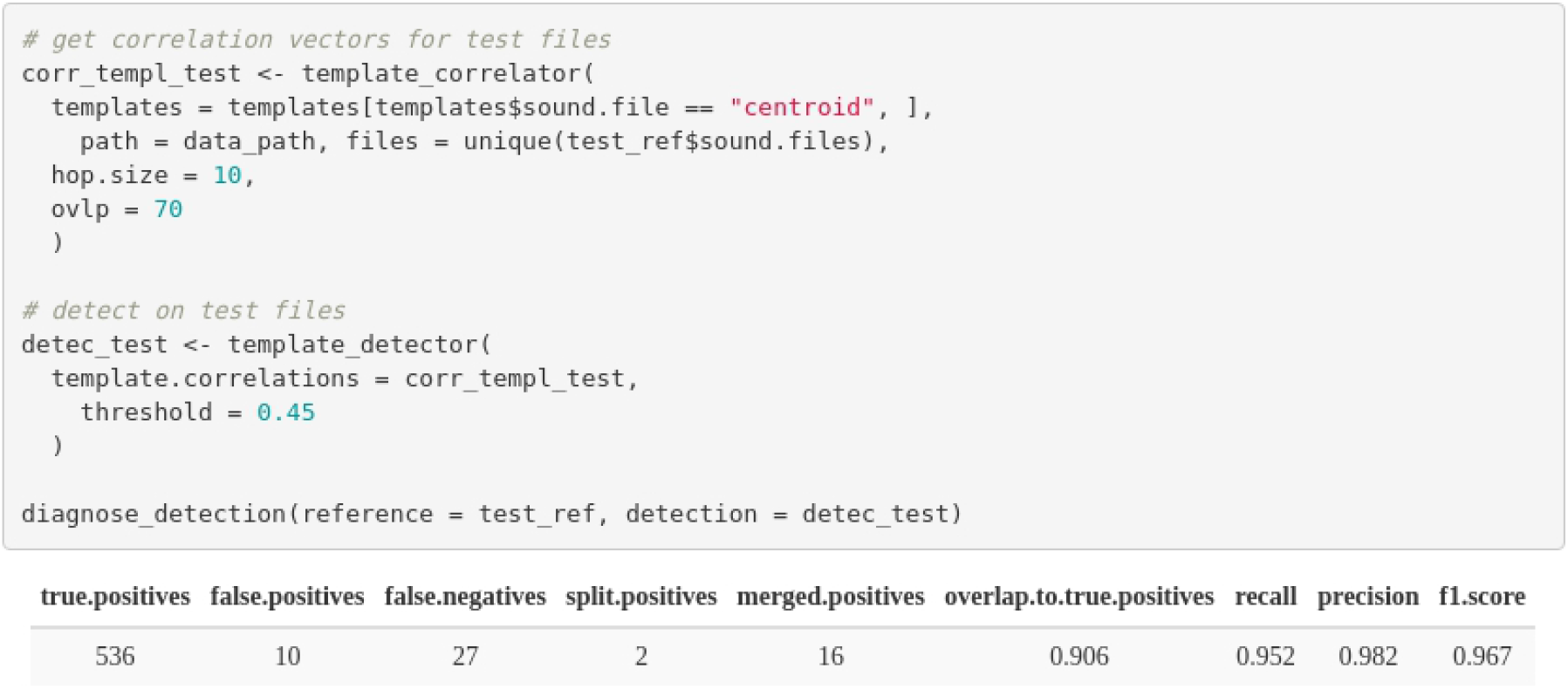

The last line of codes evaluates the detection on the test data set, which shows a good performance for both recall and precision (0.95 and 0.98 respectively).

#### Energy-based detection on zebra finch vocalizations

We used recordings from 18 zebra finch males recorded at the Rockefeller University Field Research Center Song Library (http://ofer.sci.ccny.cuny.edu/songs, Tchernichovski *et al*. 2021). Recordings contain undirected vocalizations (e.g., songs or calls) of single males recorded in sound attenuation chambers using Sound Analysis Pro. Zebra finch vocalizations are composed of multiple elements (i.e., distinct patterns of continuous traces of power spectral entropy in the spectrogram separated by time gaps) that can vary substantially in key features such as duration and frequency range (Fig. 4) and are not nearly as stereotyped as the Spix’s disc-winged bats. However, as recorded sounds show a good signal-to-noise ratio, signals in each recording can potentially be detected using an energy-based approach that does not rely on matching the acoustic structure of a template.

**Figure 4.**
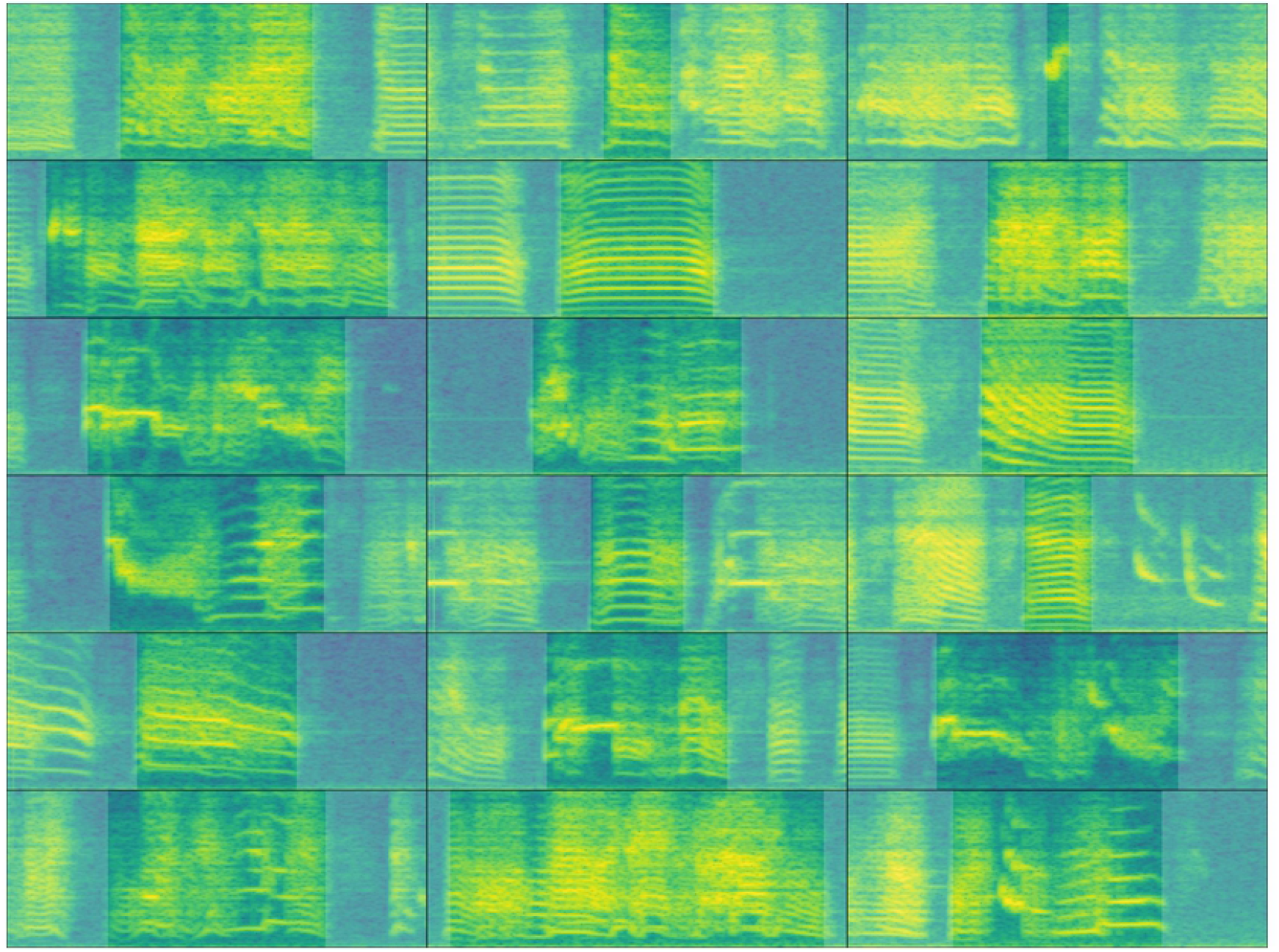
Example spectrograms of male zebra finch songs for each of the 18 sound files used in the analysis. The highest signal-to-noise ratio call by sound file are shown. The time scale range is 359 ms and the frequency range 0-11 kHz. Signals have been highlighted for visualization purposes only.

Reference annotations were made manually on the oscillogram with the spectrogram and audio as a guide using Raven Lite 2.0.1 (Cornell Lab of Ornithology). The following code loads the reference annotations and split them into two data sets for training (3 sound files) and testing (15 sound files):

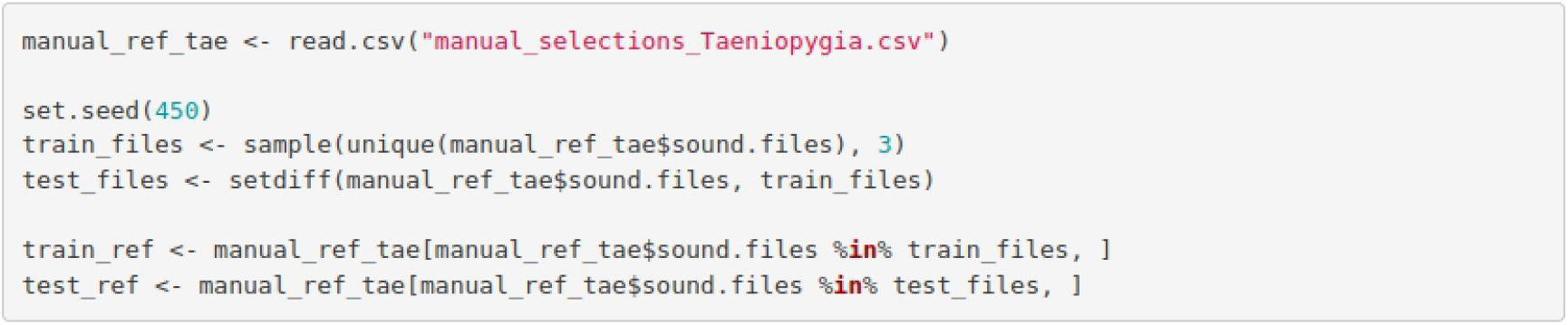

The detection parameters can be optimized using the function optimize_energy_detector. This function runs a detection for all possible combinations of tuning parameters. The code below tries three minimum duration and maximum duration values and two hold time values (which merges sounds within the specified time interval):

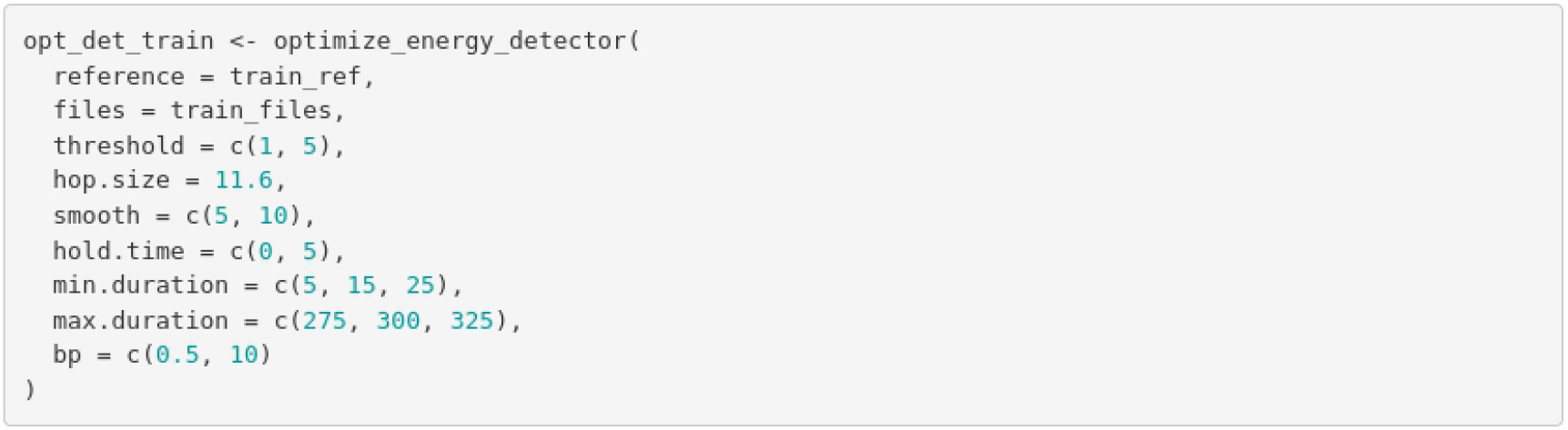

The output (opt_det_train) shows the performance indices for each of those combinations. Here we show the ten combinations with the highest F1 score:

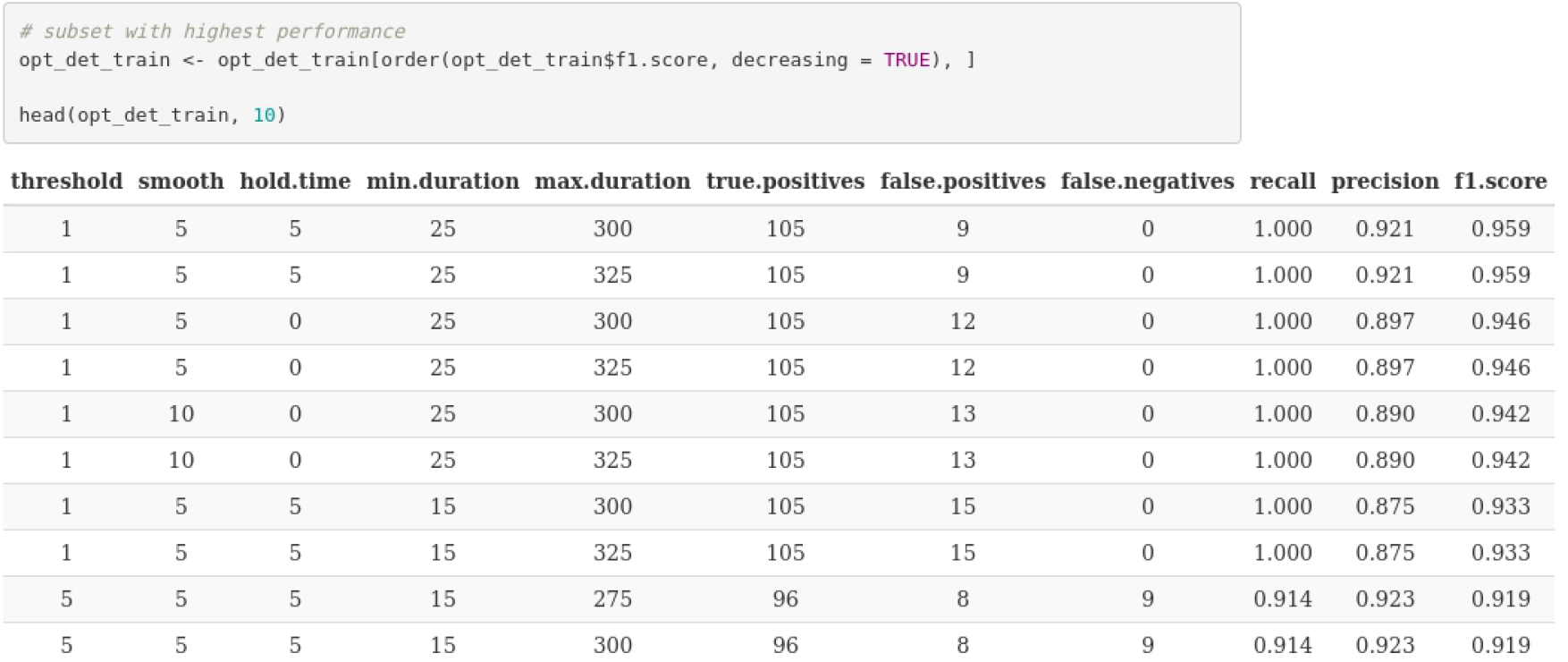

We now can use the tuning parameter values that yielded the best performance to detect sounds on the test data set:

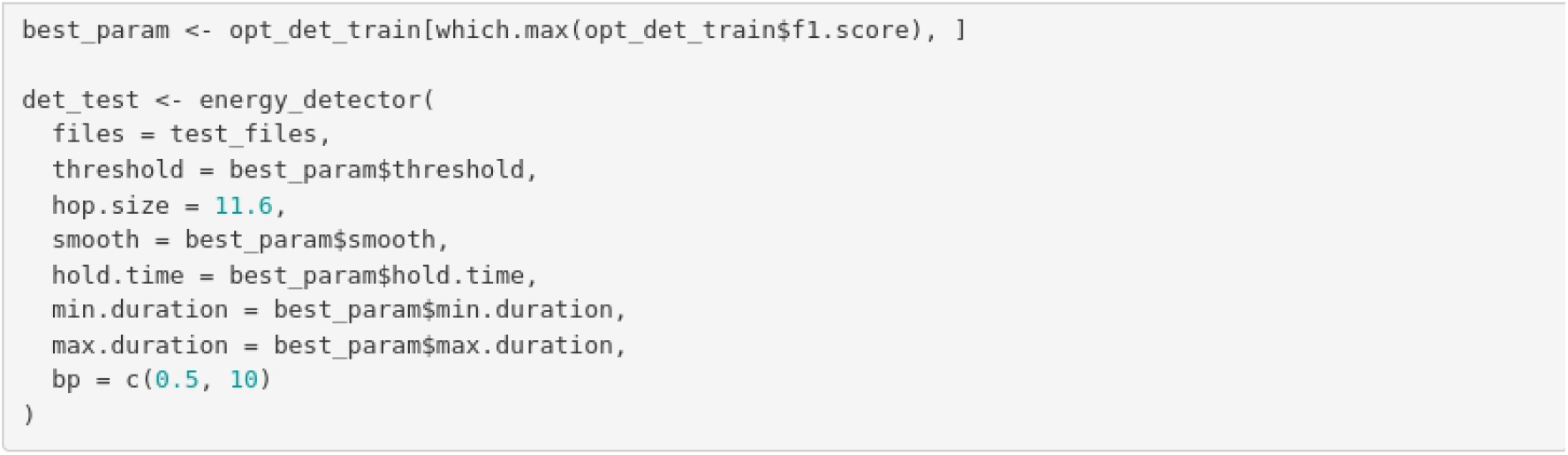

As our reference annotations include all sounds in both the training and test annotations, we can evaluate the performance of the detection on the test set as well:

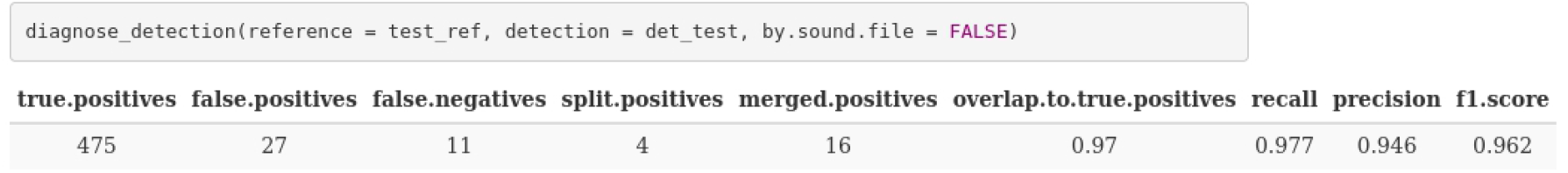

The performance on the test data set was also acceptable, with an F1 score of 0.95. Note that in the example we used a small subset of sound files for training. More training data might be needed for optimizing a detection routine on larger data sets or recordings with more variable sounds or background noise levels.

#### Additional tools

The *ohun* package offers additional tools to simplify sound event detection. Detected sounds can be labeled as false or true positives with the function ‘label_detection’. This allows users to explore the structure of false positives and figure out ways to exclude them. The function ‘filter_detections’ can remove ambiguous sounds (i.e., those labeled as split or merged detections), keeping only those that maximize a specific criterium (i.e., the highest template correlation). Finally, note that several templates representing the range of variation in signal structure can be used to detect semi-stereotyped sounds or sterepotyped multi-element repertoires when running template-based detection (‘template_detection’ function).

## Discussion

Here we have shown how to evaluate the performance of sound event detection routines using the package *ohun*. The package can evaluate detection outputs imported from other software, as well as its own detection routines. The latter can be iterated over combinations of tuning parameters to find those values that optimize detection. Although signal detection indices are commonly reported when presenting new automatic detection methods, to our knowledge, in addtion to *ohun* there is only one other performance-evaluating software developed in a free, open-source platform (sed- eval, Mesaros 2016). These two software packages can provide a common framework for evaluating sound event detection can simplify comparing the performance of different tools and selecting those tools better suited to a given research question and study system. The tools offered by *ohun* for diagnosing detection performance should not necessarily be limited to acoustic data. *ohun* can also be used for cases in which the time of occurrence of discrete events needs to be identified, such as detecting specific behaviors in video analysis of animal motor activity (*e.g*., Sturman *et al*. 2020; Hsu & Yttri 2021; Bohnslav *et al*. 2021). The detection of such motor events in video recordings can also be evaluated and optimized compared to a reference annotation, as we have shown here for sound events.

The *ohun* package provides two detection methods: template-based and energy-based detection. Compared to new deep learning approaches for finding the occurrence of sound events, the two native methods are relatively simple tools. However, these methods have been widely used by the bioacoustic community (Mellinger & Clark 2000; Charif *et al*. 2010; Aide *et al*. 2013; Specth 2002; Hafner & Katz 2015) and can reach adequate performance under the appropriate conditions, as evidenced by our two study cases and from previous reports (Knight *et al*. 2017). Deep learning methods tend to require greater computational power, larger training data sets, and, in some cases, more complex training routines (but see transfer learning approaches). This might bring unnecessary difficulties when dealing with less challenging detection tasks. Therefore, the availability of a wide range of approaches can simplify finding the most appropriate tool for the intricacies of a study system and research goals and having tools accessible to a broader research community. The tools offered in *ohun* can also be used in a subsequent pipeline in which detected sounds are further classified and false positives are mitigated using more elaborated discrimination algorithms (Balantic & Donovan 2020). Detection performance might be improved by using acoustic structure measurements to distinguish target from non-target sound events.

Detection routines can take a long time when working with large amounts of acoustic data (e.g. long recordings or many files). We provide some additional tips that can help make a routine more time efficient. 1. Always test procedures on small data subsets. Make sure to obtain decent results on a small subset of recordings before scaling up the analysis. 2. Template-based detection is almost always faster than energy-based detection. 3. Run routines in parallel. Parallelization (i.e., the ability to distribute tasks over several cores in your computer) can significantly speed up routines. All automatic detection and performance evaluation functions in *ohun* allow users to run analysis in parallel (see parallel argument in those functions). Hence, a computer with several cores can help improve efficiency. 4. Try using a computer with lots of RAM or a computer cluster for working on large amounts of data. 5. Sampling rate matters. Detecting sounds on low sampling rate files is faster, so we must avoid having Nyquist frequencies much higher than the highest frequency of the target sounds. Lastly, we underscore that these tips are not restricted to *ohun* and can also be helpful to speed up routines in other software packages.

Other things should be considered when aiming to detect sound events automatically. When running energy-based detection routines, try to use your knowledge of the signal structure to determine the initial range of tuning parameters. This can be extremely helpful for narrowing down possible parameter values. As a general rule, if human observers have difficulty detecting where a target sound occurs in a sound file, detection algorithms will likely yield low detection performance. In cases in which occurrences are ambiguous, low performances are expected. Ensure reference annotations contain all target sounds and only the target sounds. Otherwise, performance optimization can be misleading as the performance of a given detection method cannot be better than the reference itself. Lastly, avoid having overlapping sounds or several sounds as a single detection (e.g., a multi-syllable vocalization) in the reference annotation when running an energy-based detector, as they are likely to be identified as separated units.

## Supporting information

Study case Zebra Finch code

Study case Spix's disc-winged bat code

## Acknowledgments

Nazareth Rojas, Silvia Chaves-Ramírez, Mariela Sánchez-Chavarría, Miriam Gioiosa, Cristian Castillo-Salazar, and José Pablo Barrantes for their help collecting acoustic data for Spix’s disc-winged bats. We also thank the Centro Biológico Hacienda Barú for their continuous support of our research and Mijail Rojas and Andrey Sequeira for support in the early stages of package development. This study was partly funded by a CONARE-Max Planck grant from the Consejo Nacional de Rectores and the research activity C0754 at Centro de Investigación en Neurociencias, Universidad de Costa Rica. A.R-G. is supported by the Walt Halperin Endowed Professorship and the Washington Research Foundation as Distinguished Investigator.

## Ethical note

All sampling protocols followed guidelines approved by the American Society of Mammalogists for the capture, handling, and care of mammals (Sikes *et al*. 2016) and the ASAB/ABS Guidelines for the use of animals in research. This study was conducted in accordance with the ethical standards for animal welfare of the Costa Rican Ministry of Environment and Energy, Sistema Nacional de Áreas de Conservación, permit no. SINAC-ACOPAC-RES-INV-008-2017 (Decree No. 32553-MINAE). Protocols were also approved by the University of Costa Rica’s Institutional Animal Care and Use Committee (CICUA-42-2018).

## Supporting Information

Supplementary information associated with this article is available in 10.6084/m9.figshare.21675692

