## Supplementary material for "ohun: an R package for diagnosing and optimizing automatic sound event detection": Study case Zebra Finch code

study\_case\_taeniopygia.knit


### **Energy-based automatic detection on zebra finch songs**

##### **ohun**

#### 08-12-2022

##### Purpose

- Showcase energy-based automatic detection with the R package ohun
  using zebra-finch songs

##### Data set

The data used in this tutorial (annotations and recordings) can be
downloaded from here:

- https://figshare.com/ndownloader/files/38431340

##### Load package

```
library(ohun)

library(warbleR)

library(viridis)
```

##### Set directory where the sound files and annotations are found

```
data_path <- "DIRECTORY_WITH_SOUND_FILES_AND_ANNOTATIONS_HERE"
```

##### Read reference annotations

```
manual_ref <- read.csv(file.path(data_path, "manual_selections_Taeniopygia.csv"))
```

##### Create spectrograms to explore vocalization structure

This code creates a multipanel image with multiple spectrograms, one
for each of the individuals/recordings in the complete data set

```
# select highest signal to noise ratio calls per individual
manual_ref_snr <- signal_2_noise(X = manual_ref, mar = 0.05, path = data_path)

# select 1 example per sound file
high_snr <- manual_ref_snr[ave(-manual_ref_snr$SNR, manual_ref_snr$sound.files, FUN = rank) <= 1, ]

high_snr$bottom.freq <- -1
high_snr$top.freq <- 12

# create catalogs
catalog(X = high_snr, flim = c(0, 11), nrow = 6, ncol = 3, ovlp = 90, height = 15, width = 20, same.time.scale = TRUE, mar = 0.01, wl = 512, gr = FALSE, spec.mar = 0, lab.mar = 0.001, rm.axes = TRUE, by.row = TRUE, box = TRUE, pal = viridis, parallel = 10, collevels = seq(-140, 0, 5), path = data_path, img.prefix = "same.time", highlight = TRUE, alpha = 0.2)
```

##### Split reference annotations for training and testing detection

```
set.seed(450)    
train_files <- sample(unique(manual_ref$sound.files), 3)    

test_files <- setdiff(manual_ref$sound.files, train_files)

train_ref <- manual_ref[manual_ref$sound.files %in% train_files, ]
test_ref <- manual_ref[manual_ref$sound.files %in% test_files, ]
```

##### Optimize detection on training data

```
opt_det <- optimize_energy_detector(
  reference = train_ref, 
  files = train_files, 
  threshold = c(1, 5), 
  hop.size = 11.6, 
  smooth = c(5, 10), 
  hold.time = c(0, 5), 
  min.duration = c(5, 15, 25), 
  max.duration = c(275, 300, 325), 
  bp = c(0.5, 10),
  path = data_path
)
```

##### Check subset with highest performance (sorted by f1 score)

```
# subset with highest performance
sub_opt_det <- opt_det[order(opt_det$f1.score, decreasing = TRUE), ]

sub_opt_det <- sub_opt_det[1:10, c("threshold", "smooth", "hold.time", "min.duration", "max.duration", "true.positives", "false.positives",  "false.negatives", "recall", "precision", "f1.score")]

#print 
sub_opt_det
```

##### Select the highest performance parameters based on f1 score

```
best_param <- opt_det[which.max(opt_det$f1.score), ]

best_param
```

```
##    threshold peak.amplitude smooth hold.time min.duration max.duration thinning
## 45         1              0      5         5           25          300        1
##    true.positives false.positives false.negatives split.positives
## 45            105               9               0               0
##    merged.positives overlap.to.true.positives mean.duration.true.positives
## 45               12                 0.9833218                    0.1227803
##    mean.duration.false.positives mean.duration.false.negatives
## 45                     0.1158555                            NA
##    proportional.duration.true.positives duty.cycle recall precision  f1.score
## 45                              0.91706  0.3008679      1 0.9210526 0.9589041
```

##### Run detection on test data

```
det_test <- energy_detector(
  files = test_files, 
  threshold = best_param$threshold, 
  hop.size = 11.6, 
  smooth = best_param$smooth, 
  hold.time = best_param$hold.time, 
  min.duration = best_param$min.duration, 
  max.duration = best_param$max.duration, 
  bp = c(0.5, 10), 
  path = data_path
)
```

##### Evaluate detection performance

```
diagnose_detection(reference = test_ref, detection = det_test, by.sound.file = FALSE)
```

```
##   true.positives false.positives false.negatives split.positives
## 1            475              27              11               4
##   merged.positives overlap.to.true.positives    recall precision  f1.score
## 1               16                 0.9696413 0.9773663 0.9462151 0.9615385
```

##### Takeaways

- Good performance on detecting zebra finch songs: F1 score was 0.95
  for the training data set and 0.96 for the testing data

---

Session information

```
## R version 4.1.0 (2021-05-18)
## Platform: x86_64-pc-linux-gnu (64-bit)
## Running under: Ubuntu 20.04.2 LTS
## 
## Matrix products: default
## BLAS:   /usr/lib/x86_64-linux-gnu/atlas/libblas.so.3.10.3
## LAPACK: /usr/lib/x86_64-linux-gnu/atlas/liblapack.so.3.10.3
## 
## locale:
##  [1] LC_CTYPE=pt_BR.UTF-8       LC_NUMERIC=C              
##  [3] LC_TIME=es_CR.UTF-8        LC_COLLATE=pt_BR.UTF-8    
##  [5] LC_MONETARY=es_CR.UTF-8    LC_MESSAGES=pt_BR.UTF-8   
##  [7] LC_PAPER=es_CR.UTF-8       LC_NAME=C                 
##  [9] LC_ADDRESS=C               LC_TELEPHONE=C            
## [11] LC_MEASUREMENT=es_CR.UTF-8 LC_IDENTIFICATION=C       
## 
## attached base packages:
## [1] stats     graphics  grDevices utils     datasets  methods   base     
## 
## other attached packages:
##  [1] viridis_0.6.2      viridisLite_0.4.0  DT_0.20            remotes_2.4.2     
##  [5] pbapply_1.5-0      Rraven_1.0.13      ohun_0.1.0         warbleR_1.1.27    
##  [9] NatureSounds_1.0.4 knitr_1.36         seewave_2.1.8      tuneR_1.3.3.1     
## 
## loaded via a namespace (and not attached):
##  [1] xfun_0.31         bslib_0.3.1       colorspace_2.0-2  vctrs_0.3.8      
##  [5] htmltools_0.5.2   yaml_2.2.1        utf8_1.2.2        rlang_0.4.12     
##  [9] jquerylib_0.1.4   pillar_1.6.4      glue_1.5.1        lifecycle_1.0.1  
## [13] stringr_1.4.0     munsell_0.5.0     gtable_0.3.0      htmlwidgets_1.5.4
## [17] evaluate_0.14     fastmap_1.1.0     fftw_1.0-6.1      crosstalk_1.2.0  
## [21] parallel_4.1.0    fansi_0.5.0       highr_0.9         Rcpp_1.0.7       
## [25] scales_1.1.1      jsonlite_1.7.2    gridExtra_2.3     rjson_0.2.20     
## [29] ggplot2_3.3.5     digest_0.6.29     stringi_1.7.6     dtw_1.22-3       
## [33] grid_4.1.0        tools_4.1.0       bitops_1.0-7      magrittr_2.0.1   
## [37] sass_0.4.1        RCurl_1.98-1.5    proxy_0.4-26      tibble_3.1.6     
## [41] crayon_1.4.2      pkgconfig_2.0.3   MASS_7.3-54       ellipsis_0.3.2   
## [45] rmarkdown_2.14    R6_2.5.1          signal_0.7-7      compiler_4.1.0
```
