## Supplementary material for "ohun: an R package for diagnosing and optimizing automatic sound event detection": Study case Spix's disc-winged bat code

study\_case\_thyroptera.knit


### **Template-based automatic detection on Spix’s disc-winged bat social calls**

##### **ohun**

#### 13-12-2022

##### Purpose

- Showcase template-based automatic detection with the R package ohun
  using Spix’s disc-winged bat social calls

##### Data set

The data used in this tutorial (annotations and recordings) can be
downloaded from here:

- https://figshare.com/ndownloader/files/38431409

---

##### Load packages

```
library(ohun)

library(warbleR)

library(ggplot2)

library(viridis)
```

##### Set directory where the sound files and annotations are found

```
data_path <- "DIRECTORY_WITH_SOUND_FILES_AND_ANNOTATIONS_HERE"
```

##### Read reference annotations

```
manual_ref <- read.csv(file.path(data_path, "manual_annotations_thyroptera.csv"))
```

##### Create spectrograms to explore vocalization structure

This code creates a multipanel image with multiple spectrograms, one
for each of the individuals/recordings in the complete data set

```
# select highest signal to noise ratio calls per individual
manual_ref_snr <- signal_2_noise(X = manual_ref, mar = 0.05)

# select 1 example per sound file
high_snr <- manual_ref_snr[ave(-manual_ref_snr$SNR, manual_ref_snr$sound.files, FUN = rank) <= 1, ]

# create catalogs
catalog(X = high_snr, flim = c(10, 45), nrow = 6, ncol = 5, ovlp = 90, height = 10, width = 20, same.time.scale = TRUE, mar = 0.005, wl = 512, gr = FALSE, spec.mar = 0, lab.mar = 0.001, rm.axes = TRUE, by.row = TRUE, box = FALSE, pal = viridis, collevels = seq(-100, 0, 5), labels = NULL)
```

##### Split reference annotations for training and testing detection

```
set.seed(1)

train_files <- sample(unique(manual_ref$sound.files), size = 5)

test_files <- setdiff(manual_ref$sound.files, train_files)

train_ref <- manual_ref[manual_ref$sound.files %in% train_files, ]
test_ref <- manual_ref[manual_ref$sound.files %in% test_files, ]
```

##### Get diverse set of templates

```
# find templates
templates <- get_templates(train_ref, path = data_path,  bp = c(10, 50), fast = TRUE, ovlp = , wl = 200, n.sub.spaces = 3)
```

```
## The first 2 principal components explained 0.51 of the variance
```

```
# create ext. selection table
templates_est <- selection_table(templates, extended = TRUE, confirm.extended = FALSE, path = data_path)

templates_est <- rename_est_waves(templates_est, new.sound.files = templates$template)
```

##### Create spectrograms of templates

```
# create catalogs
catalog(X = templates_est, flim = c(10, 45), nrow = 2, ncol = 2, ovlp = 90, height = 10, width = 15, same.time.scale = TRUE, mar = 0.005, wl = 512, gr = FALSE, spec.mar = 0.4, lab.mar = 0.8, rm.axes = TRUE, by.row = TRUE, box = TRUE, pal = viridis, parallel = 10, collevels = seq(-100, 0, 5), img.prefix = "templates")
```

##### Extract correlation vectors and optimize correlation threshold

```
corr_templ_train <- template_correlator(
  templates = templates_est,
    path = data_path, 
  files = unique(train_ref$sound.files), 
  hop.size = 10, 
  ovlp = 70
  )

opt_detec_train <- optimize_template_detector(
  reference = train_ref, 
  template.correlations = corr_templ_train,
  threshold = seq(0.1, 0.9, 0.05)
  )
```

##### Create plot with threshold vs f1 score

##### Check subset with highest performance (sorted by f1 score)

```
opt_detec_train <- opt_detec_train[order(opt_detec_train$f1.score, decreasing = TRUE), ]

sub_opt_detec_train <- opt_detec_train[1:10, c("templates","threshold", "true.positives", "false.positives",  "false.negatives", "recall", "precision", "f1.score")]

sub_opt_detec_train
```

##### Run detection on test files

We used a threshold of 0.45 as te centroid template seems to behave
well across thresholds.

```
# get correlation vectors for test files
corr_templ_test <- template_correlator(
  templates = templates_est[templates_est$sound.file == "centroid", ],
    path = data_path, files = unique(test_ref$sound.files), 
  hop.size = 10, 
  ovlp = 70
  )

# detect on test files
detec_test <- template_detector(
  template.correlations = corr_templ_test,
    threshold = 0.45
  )
```

##### Evaluate detection performance on test files

```
diagnose_detection(reference = test_ref, detection = detec_test)
```

```
##   true.positives false.positives false.negatives split.positives
## 1            536              10              27               2
##   merged.positives overlap.to.true.positives    recall precision  f1.score
## 1               16                  0.906031 0.9520426  0.981685 0.9666366
```

```
## [1] 0.9666366
```

##### Takeaways

- Good performance on detecting Spix’s disc-winged social calls: F1
  score was 0.967 for the training data set and 0.994 for the testing
  data
- Some merged positives were found but notice that there are some
  overlapping signals in the reference annotation as well (can be checked
  with `warbleR::overlapping_sels(manual_ref)`)

---

Session information

```
## R version 4.1.0 (2021-05-18)
## Platform: x86_64-pc-linux-gnu (64-bit)
## Running under: Ubuntu 20.04.2 LTS
## 
## Matrix products: default
## BLAS:   /usr/lib/x86_64-linux-gnu/atlas/libblas.so.3.10.3
## LAPACK: /usr/lib/x86_64-linux-gnu/atlas/liblapack.so.3.10.3
## 
## locale:
##  [1] LC_CTYPE=pt_BR.UTF-8       LC_NUMERIC=C              
##  [3] LC_TIME=es_CR.UTF-8        LC_COLLATE=pt_BR.UTF-8    
##  [5] LC_MONETARY=es_CR.UTF-8    LC_MESSAGES=pt_BR.UTF-8   
##  [7] LC_PAPER=es_CR.UTF-8       LC_NAME=C                 
##  [9] LC_ADDRESS=C               LC_TELEPHONE=C            
## [11] LC_MEASUREMENT=es_CR.UTF-8 LC_IDENTIFICATION=C       
## 
## attached base packages:
## [1] stats     graphics  grDevices utils     datasets  methods   base     
## 
## other attached packages:
##  [1] DT_0.20            ohun_0.1.0         ggplot2_3.3.5      viridis_0.6.2     
##  [5] viridisLite_0.4.0  bioacoustics_0.2.7 warbleR_1.1.27     NatureSounds_1.0.4
##  [9] knitr_1.36         seewave_2.1.8      tuneR_1.3.3.1      remotes_2.4.2     
## 
## loaded via a namespace (and not attached):
##  [1] xfun_0.31         bslib_0.3.1       pbapply_1.5-0     lattice_0.20-44  
##  [5] colorspace_2.0-2  vctrs_0.3.8       htmltools_0.5.2   yaml_2.2.1       
##  [9] utf8_1.2.2        rlang_0.4.12      jquerylib_0.1.4   pillar_1.6.4     
## [13] glue_1.5.1        withr_2.4.3       sp_1.5-0          lifecycle_1.0.1  
## [17] stringr_1.4.0     munsell_0.5.0     gtable_0.3.0      moments_0.14     
## [21] htmlwidgets_1.5.4 evaluate_0.14     labeling_0.4.2    fastmap_1.1.0    
## [25] fftw_1.0-6.1      crosstalk_1.2.0   parallel_4.1.0    fansi_0.5.0      
## [29] highr_0.9         Rcpp_1.0.7        scales_1.1.1      jsonlite_1.7.2   
## [33] soundgen_2.1.0    farver_2.1.0      gridExtra_2.3     rjson_0.2.20     
## [37] digest_0.6.29     stringi_1.7.6     dtw_1.22-3        grid_4.1.0       
## [41] tools_4.1.0       bitops_1.0-7      magrittr_2.0.1    sass_0.4.1       
## [45] RCurl_1.98-1.5    proxy_0.4-26      tibble_3.1.6      crayon_1.4.2     
## [49] pkgconfig_2.0.3   MASS_7.3-54       ellipsis_0.3.2    shinyBS_0.61     
## [53] rmarkdown_2.14    R6_2.5.1          signal_0.7-7      compiler_4.1.0
```
